# Sporulation efficiency and spore quality in a human intestinal isolate of *Bacillus cereus*

**DOI:** 10.1101/2022.06.22.497182

**Authors:** Maria Vittoria, Anella Saggese, Giovanni Di Gregorio Barletta, Stefany Castaldi, Rachele Isticato, Loredana Baccigalupi, Ezio Ricca

**Affiliations:** Department of Biology, Federico II University of Naples, Italy; Department of Molecular Medicine and Medical Biotechnology, Federico II University of Naples, Italy

**Keywords:** *Bacillus cereus*, spores, temperature, biofilm, toxins

## Abstract

The *Bacillus cereus* group is a species complex of the *Bacillus* genus that includes several closely related species. Within this group, bacteria indicated as *B. cereus sensu stricto* (*B. cereus*) are the causative agent of two different types of gastrointestinal diseases associated with food poisoning. Outbreaks of this opportunistic pathogen are generally due to the resistance of its spores to heat, pH and desiccation that makes hard their complete inactivation from food products. *B. cereus* is commonly isolated from a variety of environments, including intestinal samples of infected and healthy people. We report the genomic and physiological characterization of MV19, a human intestinal strain closely related (ANI value of 98.81%) to the reference strain *B. cereus* ATCC 14579. MV19 cells were able to grow in a range of temperatures between 20 and 44°C. At the optimal temperature the sporulation process was induced very rapidly and mature spores efficiently released, however these appeared structurally and morphologically defective. At the sub-optimal growth temperature of 25°C sporulation was slow and less efficient but a high total number of fully functional spores was produced. Altogether, results reported here indicate that the reduced rapidity and efficiency of sporulation at 25°C are compensated by a high quality and quantity of released spores, suggesting the relevance of different performances at different growth conditions for the adaptation of this bacterium to diverse environmental niches.

## 1. Introduction

The *Bacillus cereus* group (*B. cereus sensu lato*) consists of several spore-forming Gram-positive bacterial species, widespread in nature as spores and vegetative cells (Liu et al., 2017; Bianco et al., 2021). The spores greatly contribute to the wide distribution of these bacteria in many environments for their resistance to extremes of temperatures and pH, to UV radiations and to the presence of lytic enzymes and toxic chemicals. The quiescent spore germinates when the environmental conditions allow cell growth and originates new vegetative cells able to grow by binary fission and eventually to sporulate again (McKenney et al., 2013).

The taxonomy of the *B. cereus* group is complex and rapidly evolving. Based on recent studies (Liu et al., 2017; Carrol et al., 2021), the group includes three well-characterized and several far less studied species. *B. cereus sensu stricto* (*B. cereus*), responsible for two types of foodborne intoxications, *B. thuringiensis*, an entomopathogen producing parasporal crystal inclusions made of proteins with insecticidal activity and *B. anthracis*, the agent of anthrax in humans and animals, are well-known species, genetically and physiologically characterized in details. The less studied species of the group include *B. mycoides* and *B. pseudomycoides*, forming rhizoidal colonies on solid media, the psychrotolerant *B. weihenstephanensis*, the animal probiotic *B. toyonensis*, the psychrotolerant and cytotoxic *B wiedmannii*, the thermotolerant and occasionally pathogen *B. cytotoxicus* and several, recently identified other species (Liu et al., 2017; Carrol et al., 2021).

*B. cereus* is a major cause of foodborne outbreaks (Glasset et al., 2016) and is responsible of two types of intoxications: the emetic gastrointestinal syndrome and the diarrheal syndrome. In addition, *B. cereus* is also involved in nosocomial non-gastrointestinal infections in immunosuppressed patients (Messelhäußer and Ehling-Schulz, 2018). These are often caused by biofilm formation on biomedical devices (Lin et al., 2022) and include septicemia, meningitis, brain abscesses, gas gangrene–like infections, pneumonia, severe ocular infections and bacteremia in preterm neonates (Jovanovic et al., 2021). The severity of the diseases caused by some *B. cereus* strains makes it important to discriminate between pathogenic and non-pathogenic strains. Such discrimination is generally based on the analysis of several genetic loci that, coding for known toxins, have been selected as markers. These include the heat-resistant, emetic toxin cereulide, synthesized by a Non-Ribosomal Protein Synthase (NRPS) encoded by the *ces* gene locus, the hemolysin BL encoded by the *hbl* genes, the non-hemolytic enterotoxin encoded by the *nhe* genes and the cytotoxin K encoded by the *cytK* gene (Owusu-Kwarteng et al., 2017; Jovanovic et al., 2021).

Toxin-producing strains of *B. cereus* are of particular concern for public health due to their ability to form biofilms and spores (Huang et al., 2020). Both biofilm and spore formation are survival strategies for *B. cereus* in facing adverse environmental conditions. The biofilm increases both the resistance properties of cells and spores and their adhesion to surfaces, therefore playing an important role in the contamination of foods, food processing and clinical equipments (Huang et al., 2020). *B. cereus* can form different types of biofilm under different growth conditions (Wijman et al., 2007) and is generally formed by proteins, carbohydrates and DNA (Wagner et al., 2009; Karunakaran and Biggs, 2011). Cells within the established biofilms can produce spores (Ryu and Beuchat, 2005; Wijman et al., 2007; Faille et al., 2014) and these have distinct properties with respect to spores derived from planktonic cells (Abee et al., 2011), appearing larger, more heat resistant and less efficient in germination (Van der Voort and Abee, 2013). The structure and the functional properties of the spore are also strongly affected by other growth and sporulation conditions such as medium composition, temperature, pH and water activity (Abee et al., 2011; Isticato et al., 2020). In *B. cereus*, that generally grows within a temperature range of 10 - 50°C with a temperature optimum between 30 and 40°C, the growth temperature influences the size and the morphology of the produced spores (Xu Zhou et al., 2017). Size and morphology, in turn, influence the functional properties of the spore with the largest ones showing the least resistance to chemicals and those surrounded by long appendages appearing more adhesive than those lacking such structures (Tauveron et al., 2006). The spore surface hydrophobicity is also affected by the growth conditions and a high hydrophobicity has been associated with a strong adhesion to surfaces (Husmark and Ro□nner, 1992; Andersson et al., 1995). In the *B. cereus* type strain ATCC 14579 the sporulation temperature acts on spore coat formation through the morphogenetic protein CotE, that is more abundantly represented in spores formed at 20°C than at 37°C (Bressuire-Isoard et al., 2016).

The aim of this study was to characterize a *B. cereus* strain isolated from a human intestinal sample. The isolate, MV19, was unambigously assigned to the *B. cereus sensu stricto* species and the presence of the genes associated with virulence and biofilm formation analyzed by whole-genome sequencing. Analyses of the growth and sporulation temperatures, the efficiency of sporulation, the yield and quality of the produced spores at different temperatures indicated that MV19 produced fully functional spores at the sub-optimal temperature of 25°C and functionally and morphologically defective spores at the optimal temperature of 42°C. These results suggest that producing different spores at different growth conditions is an important feature for the adaptation of spore formers to diverse environments.

## 2. Materials and Methods

### 2.1 Bacterial isolation, spore production and purification

Faecal samples were collected from healthy children not under antibiotic or probiotic treatment in the frame of a project aimed at collecting spore formers of the *Bacillus* genus of intestinal origin. About 1 g of a sample from a 4-year-old child was homogenized in PBS and heat-treated for 20 minutes at 80°C, serially diluted, plated on Difco sporulation medium (DSM) (for 1L: 8 g/L Nutrient Broth, 1 g/L KCl, 1 mM MgSO_4_, 1 mM Ca(NO_3_)_2_, 10 μM MnCl_2_, 1 μM FeSO_4_, Sigma-Aldrich, Germany) and incubated overnight at 37°C, under aerobic conditions. The selected colonies were re-streaked on DSM agar plates to isolate single colonies that were then checked for the formation of phase-bright spores under the light microscope. Spores were prepared at the required temperature by the exhaustion method (Nicholson and Setlow, 1990) using liquid DSM. Before purification, spores were washed four times with cold sterile distilled water and centrifuged at 8000 g for 20 minutes and then purified with ethanol as described by Zhao et al., 2008.

### 2.2 Physiological properties of the isolated strain

Hemolytic activity was assessed by spotting 5 μL of growing cells on Columbia agar plates supplemented with 5% defibrinated horse blood (Thermo Scientific), incubated for 24h at 37°C and analyzed for the presence of a lysis halo around the colony.

Biofilm formation was tested by growing bacterial cells in 24-well culture plates. The experiment was performed either in LB medium (10 g Bacto-Tryptone, 5g Bacto-yeast extract, 10g NaCl, pH 7.0), or in S7 minimal medium (50 mM morpholine-propane-sulfonic acid (MOPS) (adjusted to pH 7.0 with KOH), 10 mM (NH_4_)_2_SO_4_, 5 mM potassium phosphate (pH 7.0), 2 mM MgCl_2_, 0.9 mM CaCl_2_, 50 μM MnCl_2_, 5 μM FeCl_3_, 10 μM ZnCl_2_, 2 μM thiamine hydrochloride, 20 mM sodium glutamate, 1% glucose, 0.1 mg/mL phenylalanine, and 0.1 mg/mL tryptophan) or in DSM, for 48 h without shaking, at 25 and 42°C. Then, biofilm production was assessed by crystal violet assay as previously described in Petrillo et al., 2021. Data were normalized by total growth estimated by OD_590nm_, and the experiment was performed in triplicate.

The growth rate at different temperatures (25, 37 and 42°C) was measured by OD_590nm_ readings of cells growing in LB medium.

### 2.3 Physiological properties of MV19 spores

Spore resistance to chemical and physical treatments was tested as previously reported by Bressuire-Isoard et al. 2016, with some modifications. Lysozyme resistance was tested by following changes in CFU counts on LB agar plates after 1 hour of incubation of the spores (1.0×10^7^ CFU ml-1) with Lysozyme (7U mL-1) at 37°C.

Hydrogen peroxide (H_2_O_2_) resistance was tested by following changes in CFU counts on LB agar plates after 7.5 and 15 minutes of incubation of heat-activated (10 min at 70°C) spore suspension (1.0×10^7^ CFU ml-1) in a 5% H_2_O_2_ solution.

Similarly, 0.2 mL of heat-activated spores (1.0×10^7^ CFU ml-1) were incubated at 90°C for 15 and 30 minutes and then spores resistance to heat treatment was measured by following changes in CFU counts on LB agar plates.

Spore hydrophobicity was assessed by mixing 1.9-ml volume of spore suspension (0.4-0.5 OD_590nm_) in water with 0.1 ml of *n*-hexadecane and following changes in optical density at 600 nm after separation of the spores between the aqueous and apolar phases as previously reported by Bressuire-Isoard et al. 2016. The ability of spores to form clumps was evaluated as described by Cangiano et al. 2014.

Two methods were used to evaluate spore germination. Spores (1.0×10^7^ CFU ml-1) activated at 70°C for 20 minutes, were resuspended in LB medium, serially diluted and plated on LB agar plates for CFU counts. Then, the germination efficiency was assessed by flow cytometry as previously described in Cangiano et al. 2014. Heat-activated spores were stained with 0.5 μM Syto 16 (Life Technologies, Waltham, MA) and incubated in the dark for 15 min at 30°C. λ max values for the absorption and fluorescence emissions of the complex with DNA are 488 and 518 nm, respectively. Then, spores were analysed in a flow cytometer (BD Accuri C6; BD Biosciences, San Jose, CA) and detected in FL1 by using the standard C6 filter configuration (FL1 =530/30 BP) separating germination-unspecific and germination-specific fluorescence (threshold 10^4^) (Petrillo et al. 2020).

The sporulation efficiency was measured by growing cells in DSM 25 and 42°C and counting at various time points the total number of cells, sporangia, and spores under the phase-contrast microscope and measuring OD_600nm_ at spectrophotometer. All experiments were carried out in triplicate from independently purified spores.

### 2.4 Whole-genome sequencing and phylogenetic analysis

Exponentially growing cells were used to extract chromosomal DNA as previously reported (Cutting and Vander Horn, 1990). Genome sequencing of MV19 genome was performed by GenProbio (Parma, Italy) with Illumina MiSeq Sequencing System. Genome assembly was performed with SPAdes v3.14.0 by means of MEGAnnotator pipeline (Lugli et al., 2016). Open reading frames (ORFs) prediction was performed with RAPSearch2 against NCBI RefSeq database and HMMER against PFAM database. Ribosomal RNA genes prediction was performed with RNAmmer v1.2. Transfer RNA gene prediction was performed with tRNAscan-SE v1.21. The 16S rRNA gene of the isolated strain was extracted from the sequenced genome and was compared to 22 reference 16S rRNA genes of closely related strains identified using the Genome Taxonomy Database (GTDB) taxonomy and retrieved from the NCBI database. All sequences were aligned using Seaview 5.0.5 software (Petrillo et al., 2021) and the phylogenetic tree was created using the Maximum-likelihood algorithm with model GTR+I+G4. Statistical support was evaluated by the approximate likelihood-ratio test (aLRT) and is indicated at each node in the tree. *Bacillus subtilis* subsp. *subtilis* 168 was used as an outgroup to root the tree. Average Nucleotide Identity (ANI) value between the sequenced genome and the closest bacteria was obtained using GTDB-Tk Classify - v1.6.0 toolkit (Chaumeil et al., 2019). The genome of the isolated strain has been deposited in GenBank as BioProject PRJNA846192 whose accession number is CP098734.

### 2.5 Whole-genome typing of MV19

The presence of putative toxins and virulence factor-associated genes was evaluated by analyzing the MV19 genome by using the VF analyzer pipeline via a comparison to the Virulence Factor Database (VFDB, Liu et al., 2019). Nucleotide sequences of genes involved in biofilm formation and the proteins involved in the formation of coat and exosporium were found in NCBI program (https://www.ncbi.nlm.nih.gov/) and the alignments with the MV19 genome were performed using BLASTN and BLASTP, respectively.

### 2.6 Microscopy analyses

Five microliters of freshly purified spores were spotted on microscope slides and covered with coverslips and observed under phase-contrast light microscope by using an Olympus BX51 microscope fitted with a 100× objective UPlanF1. Images were captured using an Olympus DP70 digital camera equipped with Olympus U-CA Magnification Changer.

For SEM analysis, MV19 spores (1.0×10^8^/mL) prepared at different temperatures were cut from subapical parts using a sharp razor blade, fixed with 3% glutaraldehyde in phosphate buffer (65 mM, pH 7.2–7.4) for 2 h at room temperature, post-fixed with 1% osmium tetroxide in the same phosphate buffer for 1.5 h at room temperature, and completely dehydrated with ethanol. Both samples were then mounted on aluminium stubs, coated with a thin gold film using an EdwardE306 Evaporator, and observed with a FEI (Hills-boro, OR, USA) Quanta 200 ESEM in high vacuum mode (P 70 Pa, HV 30 kV, WD10 mm, spot 3.0) (Saggese et al. 2022).

## 3. Results and Discussion

### 3.1 Phylogenetic analysis and functional annotations of MV19 genome

The general genomic features of MV19 were summarized in Table 1 and Fig. 1. MV19 was initially assigned to the *Bacillus cereus* group by 16S rRNA gene sequence alignment and a phylogenetic tree constructed by comparison with 22 type strains representative of the *Bacillus cereus* group (Fig. 2). Since the sequence of the 16S gene is often not sufficient to allow a correct species assignment (Liu et al, 2015), whole genome comparison was performed (by GTDB-Tk Classify - v1.6.0), and an Average Nucleotide Identity (ANI) value of 98.81% was measured with its closest relative, the *Bacillus cereus* ATCC 14579 strain, therefore, allowing an unambiguous taxonomic classification as *Bacillus cereus sensu stricto* (Carrol et al., 2021).

**Table 1.** Genome properties of MV19

| Species | Size (bp) | Contigs | GC content (%) | Predicted ORFs |
| --- | --- | --- | --- | --- |
| *Bacillus cereus* | 5.792.430 | 98 | 35 | 5803 |

**Fig. 1:**
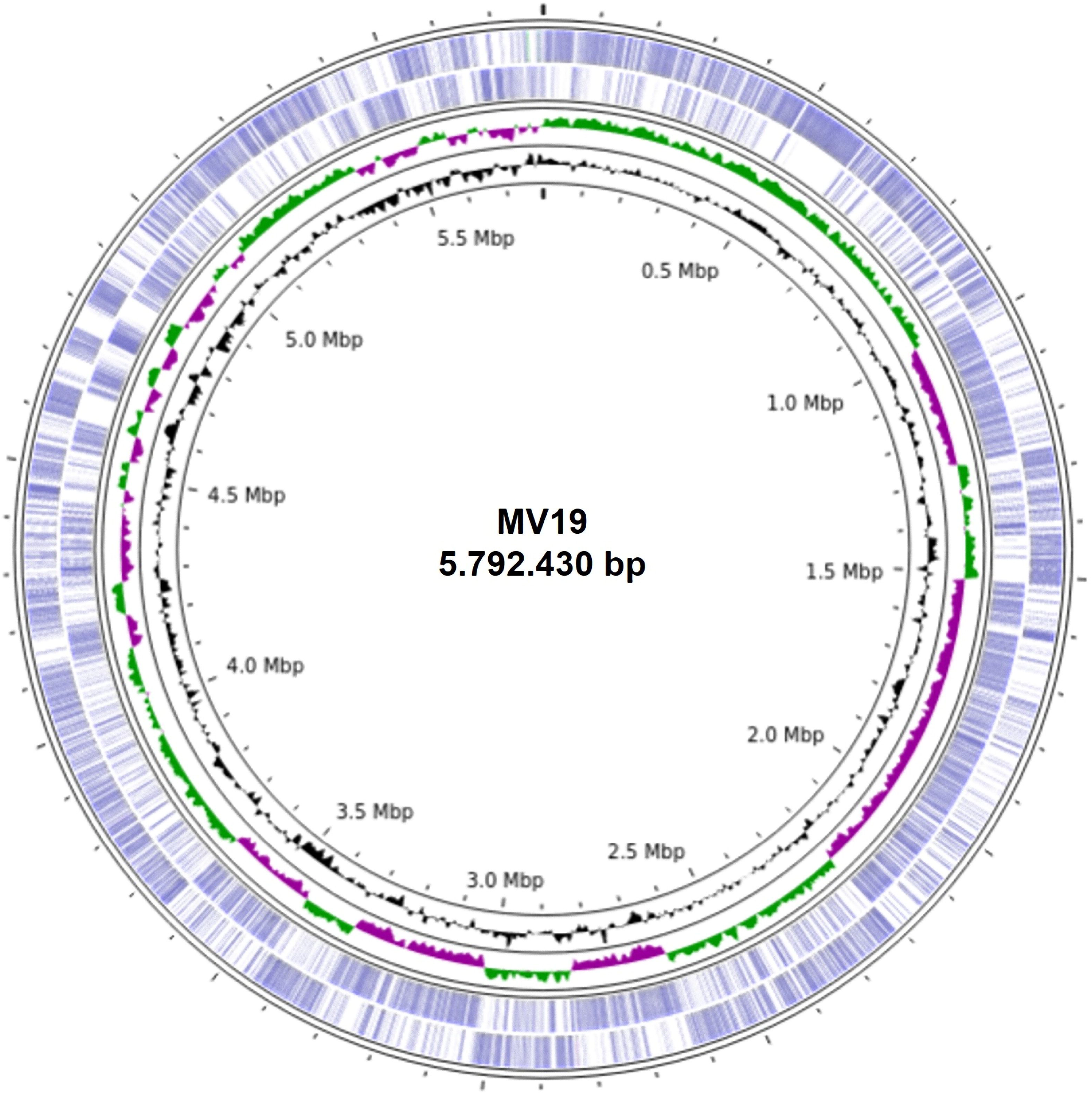
The circles represent from inside to outside: circle 1, DNA base position (bp); circle 2 and 3, GC content and GC skew; circle 4 protein-coding regions transcribed on the plus strand (clockwise); circle 5, protein-coding regions transcribed on the minus strand (anticlockwise). The genome plot was generated using Proksee.

**Fig. 2:**
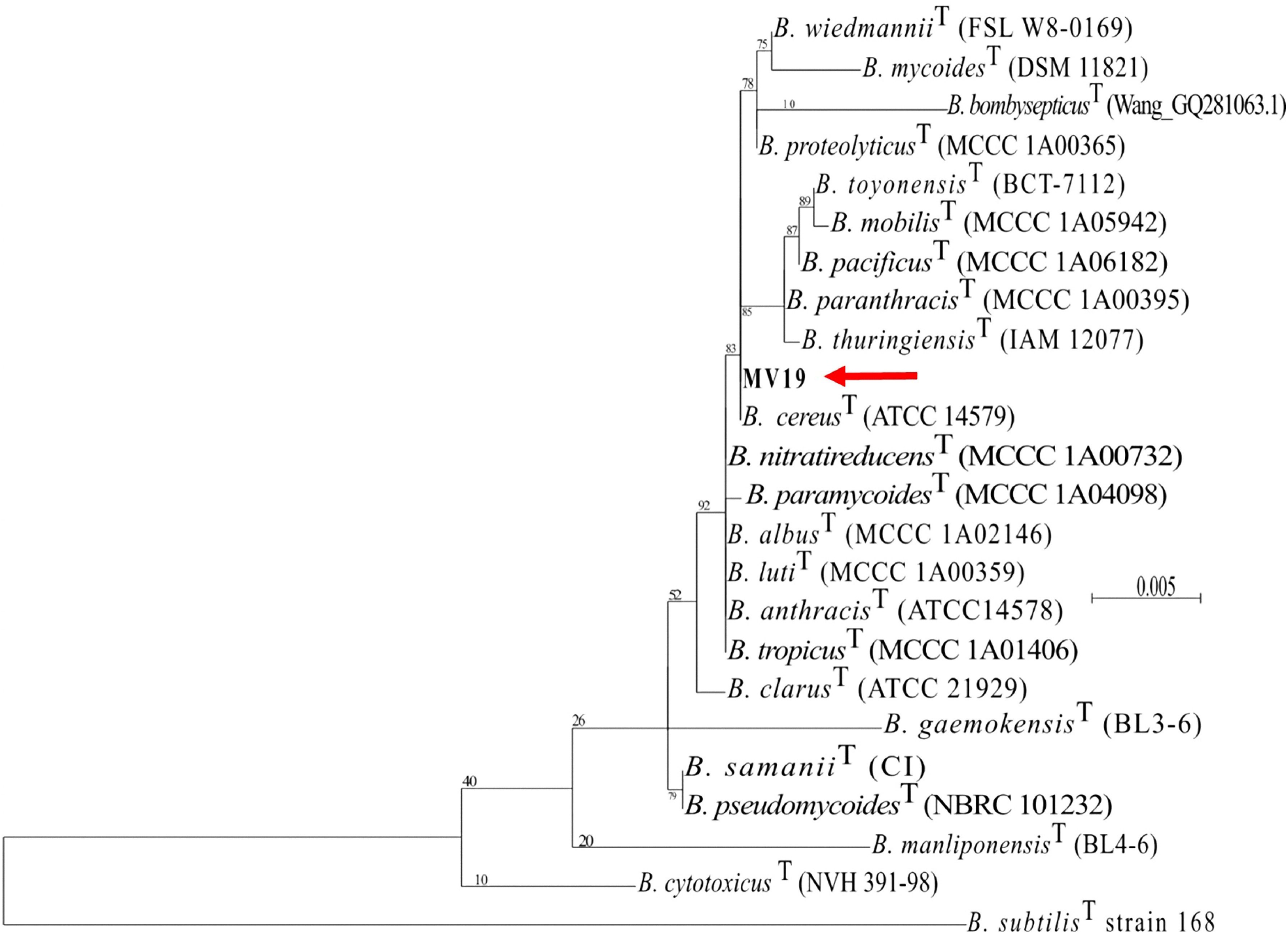
The phylogenetic tree was constructed using the Maximum likelihood algorithm with model GTR + I + G4, based on 16S rRNA gene sequences. The genome sequence of MV19 was aligned to the type strains of the *“Bacillus cereus* group” according to Genome Taxonomy Database (GTDB). Node support represents the approximate likelihood-ratio test (aLRT) and is shown at the corresponding node of the tree. *Bacillus subtilis* strain 168 is used as an outgroup.

Functional annotation of the MV19 genome was performed using the KO (KEGG Orthology) database using BlastKOALA tools (available at KEGG Web site https://www.kegg.jp/). A total of 5814 protein-coding sequences, 11 rRNAs, 68 tRNAs and 1 tmRNA were assigned. As reported in Suppl. Mat. Fig. S1A Kegg analysis showed that only 45% of the annotated genes could be assigned to subsystems. Among the subsystem categories present in the genome, “genetic information processing” and “signaling and cellular processing” had the highest feature counts, followed by “carbohydrate metabolism”. The other less represented categories are reported in Suppl. Mat. Fig. S1B with their relative percentages. MV19 genome was further classified into orthologous groups based on their function (COG), by using eggNOG (http://eggnog-mapper.embl.de/) (Suppl. Mat. Fig. S2). The majority of the annotated genes (1353 genes) were classified into “Function unknown” (S category in Suppl. Mat. Fig. S2), the next most represented class was “Amino acid transport and metabolism” (E category, 467 genes) followed by “Transcription” (K category, 451 genes).

### 3.2 Genome-wide detection of toxin genes and other virulence factors in MV19

Pathogenic strains of *B. cereus* are characterized by the presence of genes coding for toxins and other virulence-related factors (Owusu-Kwarteng et al., 2017; Jovanovic et al., 2021). To evaluate their putative presence in MV19, the genome was analyzed with the VFAnalyzer tool (http://www.mgc.ac.cn/VFs/), based on the VFDB database. As shown in Suppl. Mat. Table S1, MV19 genome contains genes coding for the Non-hemolytic enterotoxin (*nheA, nheB, nheC*), the hemolytic enterotoxin (*hblA, hblC, hblD*), the single protein cytotoxin K (*cytK*) and the hemolysin Cereolysin O (*alo*). In addition, it contains genes for the Hemolysin III (*hlyIII*) and for a protein belonging to the Hemolysin-III family proteins, while a gene coding for the Hemolysin II (*hlyII*) was not found. MV19 genome contains only one of the genes required for Cereulide synthesis, *cesH* (Suppl. Mat. Table S1). The Cereulide biosynthetic genes are organized in a cluster of seven genes (*cesH, cesP, cesT, cesA, cesB, cesC*, and *cesD*) that are located on a megaplasmid. Six genes form an operon (*cesPTABCD*) while the seventh (*cesH*) is transcribed by its own promoter and is located at the 5’ end of the operon in the same reading frame (Lücking et al., 2015). In MV19 the *cesPTABCD* operon is not present while the single gene present (*cesH*) is found on the chromosome.

In addition, genes coding for virulence-related factors such as phospholipases, sphingomyelinases, proteinases and peptidases were also found in the MV19 genome (Suppl. Mat. Table S2). In particular, the MV19 genome contains two ORFs encoding the Immune Inhibitor A metalloproteinase (*inhA*), needed to escape the immune surveillance and promote germination inside the host (Enosi Tuipulotu et al., 2021), the Phosphatidylcholine-preferring phospholipase C (*plcA*), the Phosphatidylinositol-specific phospholipase C (*piplc*) and the Sphingomyelinase (*sph*), all involved in mammalian membrane-damaging (Suppl. Mat. Table S2). A gene coding for a homolog of the Streptococcal enolase (*eno*) was also found in the genome sequence of MV19 (Suppl. Mat. Table S2). This is a secreted protein also found in different *B. anthracis* strains, where it may contribute to pathogenicity by binding the infected host’s plasminogen (Lamonica *et al,* 2005).

The ability to acquire iron is also considered a virulence-related factor, as it is crucial for bacterial growth and colonization of host tissues (Segond et al, 2014). Genes coding for two important siderophores useful for iron acquisition from ferritin, bacillibactin (*dhbABCEF*) and petrobactin (*asbABCDEF*), and the Iron-regulated leucine rich surface protein type A (*ilsA*), that acts as ferritin surface receptor, were all found in the MV19 genome (Suppl. Mat. Table S2).

Regulation of virulence gene expression occurs through a precise regulatory circuit, which in *B. cereus* is provided by a quorum sensing mechanism controlled by the transcriptional factor PlcR and the signalling peptide PapR (Ehling-Schulz et al., 2018). As shown in Suppl. Mat. Table S2, genes coding for their homologs are present in MV19 genome.

Altogether, MV19 contains genes for most known virulence factors of pathogenic *B. cereus* and should therefore be considered as a potentially virulent strain.

### 3.3 Physiological characterization

MV19 was isolated from a faecal sample of a healthy, four-year old boy not under antibiotic or probiotic treatment. The sample was heat-treated at 80°C for 20 minutes to eliminate all vegetative cells, plated on DSM and incubated aerobically to isolate *Bacillus* spore formers. Purified colonies appeared after 16-18 hours of incubation at 37°C and appeared creamy, mucoid and of irregular shape. Under the light microscope cells of MV19 were rod shaped and formed phase bright, ellipsoidal spores localized in central or sub-terminal position (respectively white or yellow arrows in Fig. 3A). In addition to spores, some MV19 cells also produced one or two phase bright structures per cell (red arrows in Fig. 3B). Production of such structures, resembling inclusion granules, was independent on the growth medium, being produced in rich (LB), minimal (S7) or sporulation-inducing (DS) media, and on the growth phase (not shown). Some forming spores (Fig. 3C) and free spore (white arrows in Fig. 3D) showed an unusual curved morphology (see below).

**Fig.3:**
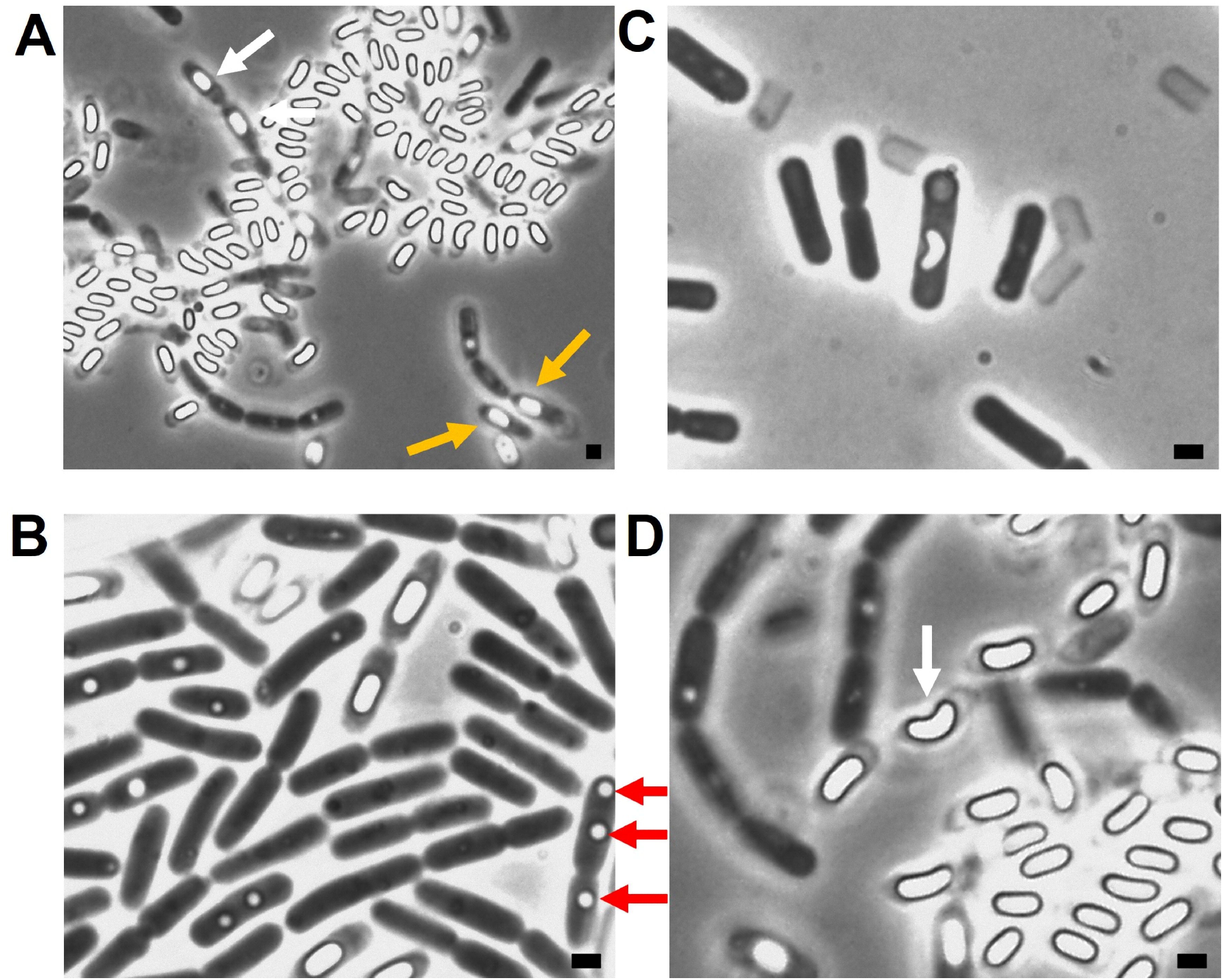
Cells of MV19 grown on solid DSM at 42°C and observed under the phase-contrast microscope. (A) Within sporangia spores have central (white arrows) or sub-terminal (yellow arrows) position. (B) Some cells contain phase bright structures (red arrows) resembling inclusion granules. Some forming spores (C) and free spore (D, white arrow) showed an unusual curved morphology. Scale bars correspond to 1 μm.

A panel of relevant antibiotics was used to assess the susceptibility of MV19 cells. As reported in Table 2, MV19 cells showed MIC for vancomycin higher than the breakpoint levels indicated for members of the *Bacillus* genus (Saggese et al. 2021) and a weak resistance to streptomycyn and chloramphenicol (Table 2).

**Table 2.** Minimal Inhibitory Concentration (MIC)^a^ of selected antibiotics against MV19

| Antibiotic | *B. cereus* MV19<sup>b</sup> | *Bacillus* EFSA breakpoint<sup>b</sup> |
| --- | --- | --- |
| Vancomycin | 12.00 | 4 |
| Clindamycin | 0.25 | 4 |
| Gentamicin | 0.38 | 4 |
| Erythromycin | 0.38 | 4 |
| Streptomycin | 2.00 | 8 |
| Tetracycline | 0.25 | 8 |
| Chloramphenicol | 4.00 | 8 |
| Kanamycin | 1.50 | 8 |

<sup>a</sup> Analysis performed with paper strips with a concentration range of 0.016-256 µg mL<sup>-1</sup> for all the antibiotics tested; <sup>b</sup> values in µg mL<sup>-1</sup>.

MV19 cells were able to completely lyse red blood cells on plate and were, therefore, indicated as β-hemolytic (not shown). Cells were able to grow in a temperature range between 20 and 44°C (not shown) and produced biofilm both on plate and on liquid cultures. The amount of biofilm formed was higher minimal (S7) than in rich (LB) or sporulation-inducing (DS) medium after 48 hours of growth (Fig. 4). Genes putatively involved in biofilm formation were identified in the MV19 genome (Suppl. Mat. Table S3). In particular, the *pur* genes, required for purine biosynthesis, the *eps* genes, coding for the exopolysaccharide, the *tapA* gene needed for biofilm assembly (Vlamakis et al., 2013 and Yan et al., 2017), and the *sinR* gene coding for the master regulator of biofilm formation (Yan et al., 2017) were all present in the MV19 genome (Suppl. Mat. Table S3).

**Fig. 4:**
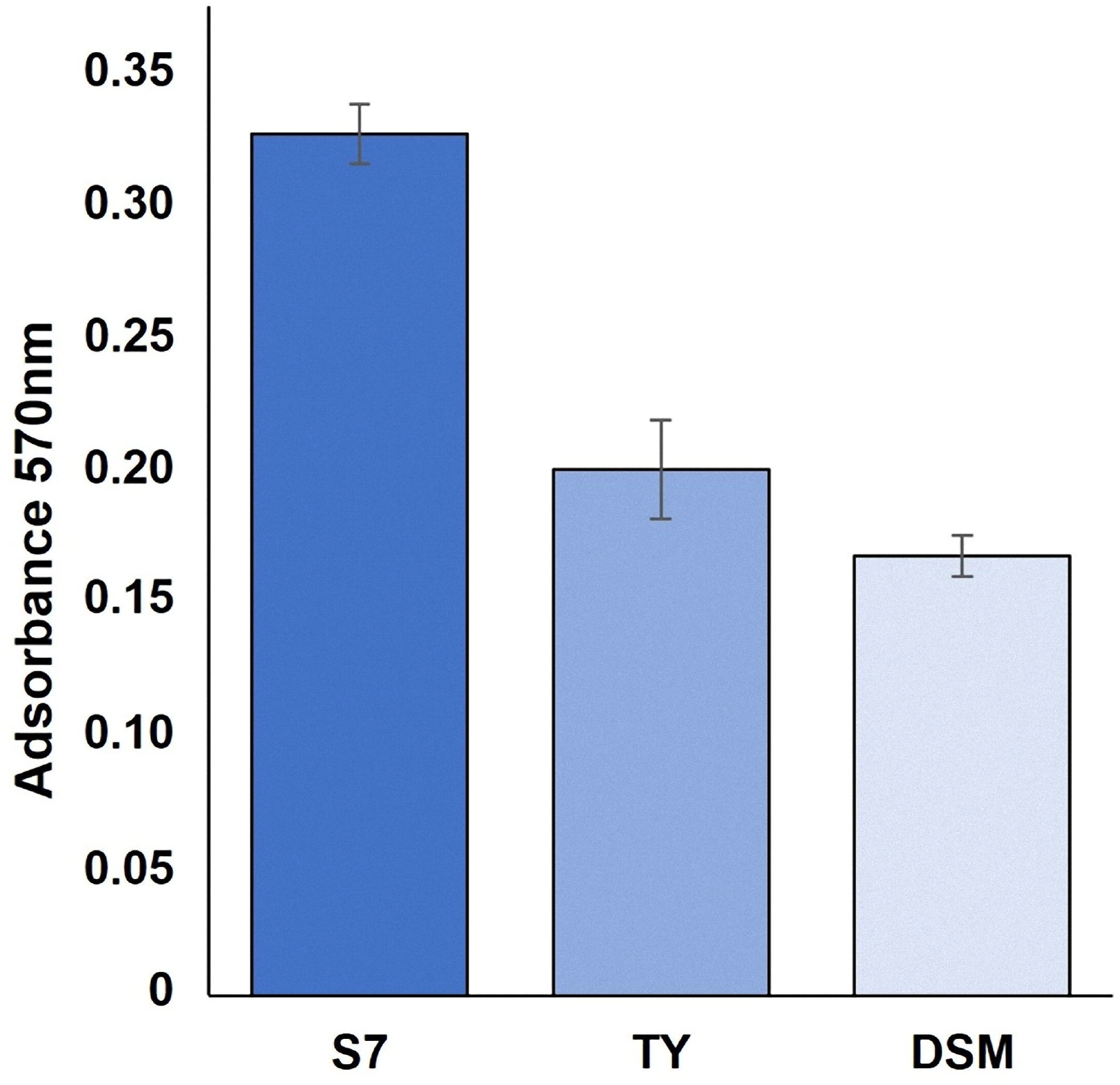
Biofilm production in minimal (S7), rich (LB) and sporulation-inducing (DS) medium at 25°C. The graph shows the OD_570nm_ values of the samples stained with crystal violet. The graph reports the average of three independent experiments.

Cell growth was followed at 25, 37 and 42°C in LB medium. At 37 and 42°C the generation time was identical (33 min) while at 25°C was longer (50 min). Moreover, at 37 and 42°C cells very rapidly started their exponential growth and quickly entered into the stationary phase of growth. At 25°C cells experienced a long lag phase, independently from the temperature used to obtain the starting inoculum, before entering into the exponential phase of growth. Therefore, 37 and 42°C were considered as optimal growth temperatures and 25°C as a sub-optimal condition in LB medium. Based on this, all further experiments were performed at the optimal temperature of 42°C and at the sub-optimal temperature of 25°C.

### 3.4 Sporulation and spore properties

MV19 were grown in DSM to induce spore formation. As in LB also in DSM cells grew more rapidly at 42°C than at 25°C (Fig. 5A). At the latter temperature entry into the exponential phase of growth was preceded by a long lag phase and a precise initiation of the sporulation cycle (T0) was not clearly identified. Indeed, after 18 hours of growth at 25°C, during the exponential phase of growth, a large number of sporulating cells was already present (see below).

**Fig.5:**
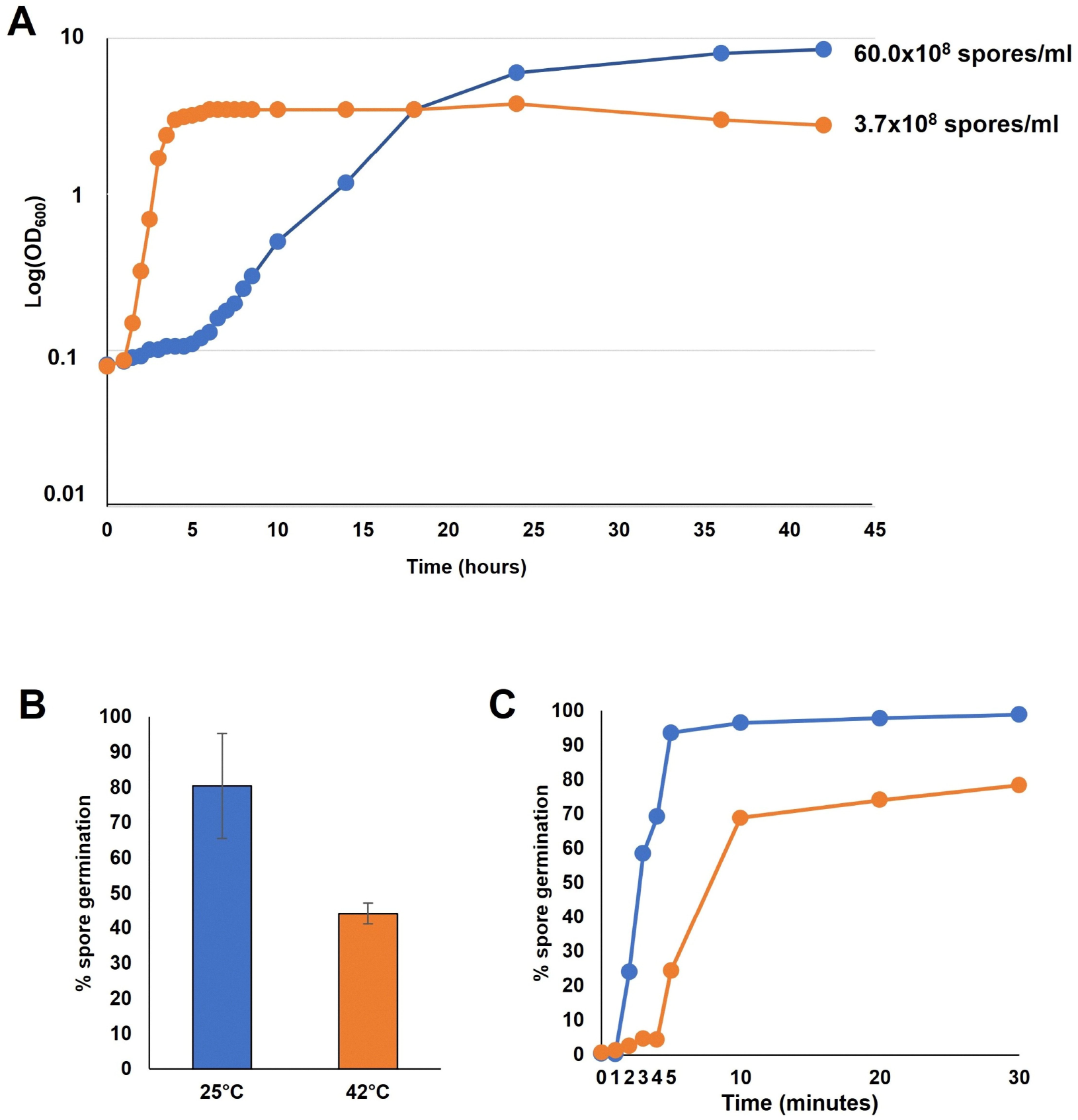
MV19 growth at 25°C (blue) or 42°C (orange) in sporulation-inducing medium (A). Germination efficiency measured by CFU counts (B) or flow cytometry (C) with spores produced at 25 (blue) and 42 °C (orange). Panel C reports the percentage of germination based on the flow cytometry data (Supplementary material Fig. S3).

Spore formation was followed by collecting samples from the growth cultures at various times and counting vegetative cells, sporangia and free spores under the light microscope. As reported in Table 3, at 42°C few free spores (5%) were produced after 18 hours of growth. These became 15% after 24 hours and after 42 hours only free spores were observed in the culture. At 25°C no free spores were observed after 18 and 24 hours and after 42 hours about 25% of sporangia were still present (Table 3). Spore production occurred more rapidly and efficiently at 42°C than at 25°C but at 25°C an over 15-fold higher number of spores was obtained (3.7×10^8^ and 60.0×10^8^ spores/ml at 25 and 42°C, respectively) (Fig. 5A). As previously reported for *B. subtilis,* a higher number of spores produced by slow-than fast-growing cells is due to a prolonged growth that allows for additional cell divisions (Mutlu et al., 2018) as also occurs for MV19 (Fig. 5A).

**Table 3.** Efficiency of spore formation at 25 and 42°C^a^.

| Temperature | Hours | Vegetative cells (%) | Sporangia (%) | Free spores (%) |
| --- | --- | --- | --- | --- |
| **25°C** | 0 | 100 | - | - |
|  | 18 | 50 | 50 | - |
|  | 24 | - | 100 | - |
|  | 42 | - | 25 | 75 |
| **42°C** | 0 | 100 | - | - |
|  | 18 | 20 | 75 | 5 |
|  | 24 | 15 | 70 | 15 |
|  | 42 | - | - | 100 |

<sup>a</sup>Cells were grown aerobically in DS medium. At the indicated times aliquots were collected and analyzed under the light microscope. The % of vegetative cells, sporangia and free spores was calculated by counting over 300 bacteria in different microscopy fields.

Spores produced at 25 or 42°C were then evaluated for their resistance properties and the germination efficiency. The efficiency of germination was assessed by plate count after suspension of purified spores in LB medium (Fig. 5B) and by flow cytometry (Fig. 5C and Suppl. Mat. Fig. S3), as previously described (Black et al., 2005; Cangiano et al., 2014). The two approaches gave similar results (Fig. 5BC) and indicated that spores produced at the sub-optimal temperature of 25°C germinated more rapidly and efficiently than those produced at the optimal growth temperature.

Spores produced at the sub-optimal temperature were also more resistant than those produced at 42°C to lysozyme (Fig. 6A), H_2_O_2_ and heat (Fig. 6B) and were more hydrophobic (Fig. 6C). Due to the high hydrophobicity of 25°C spores, when suspended in water they tended to form clumps that precipitated to the bottom of the cuvette causing a decrease of OD_590nm_ readings over time (Fig. 6D). Such effect was not observed when spores were resuspended in 50% methanol (not shown), confirming that it was due to the spore hydrophobicity.

**Fig. 6.**
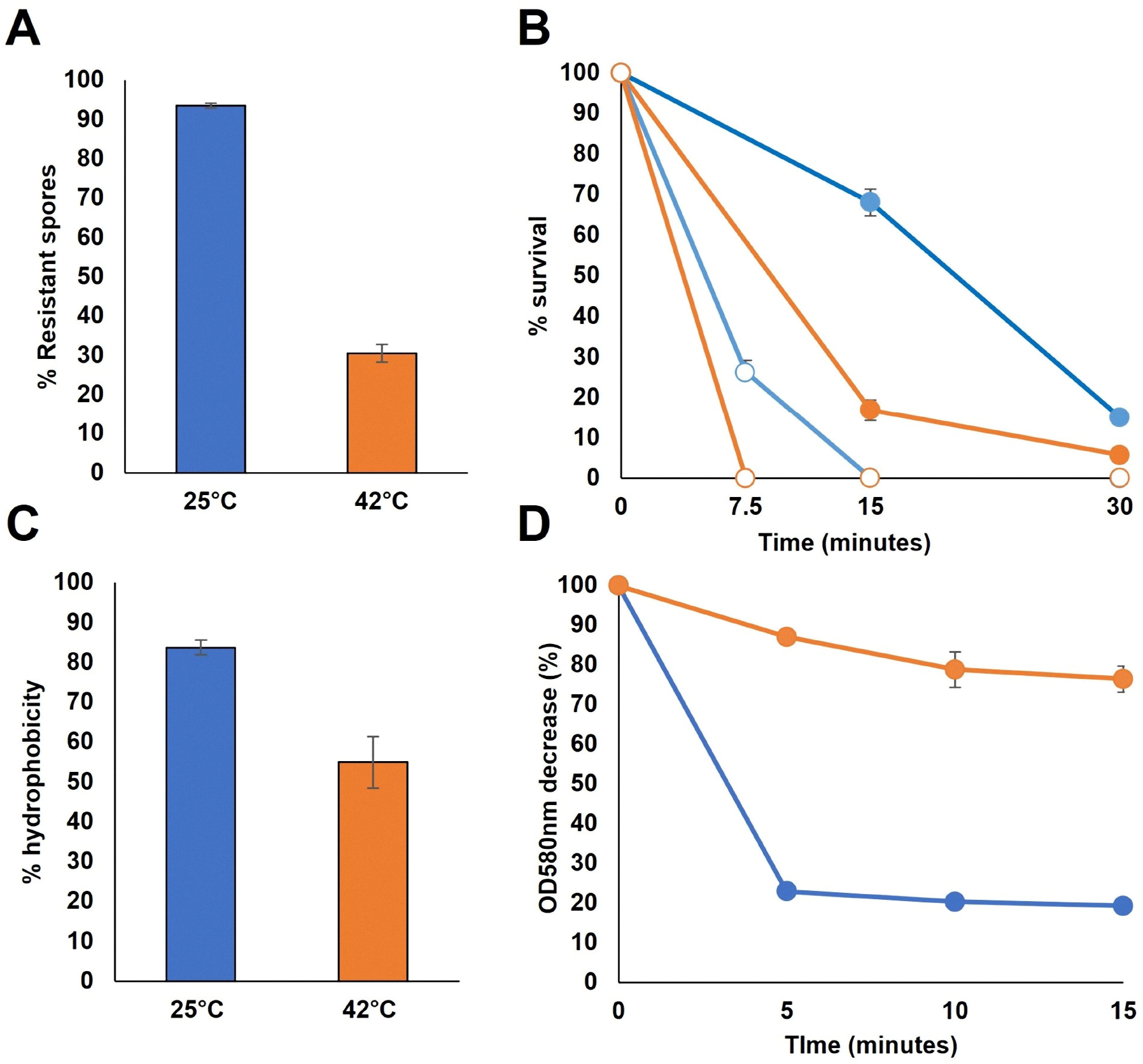
Physiological properties of MV19 spores produced at 25 (blue) and 42°C (orange). A) Lysozyme resistance assay. Percentage of resistant spores after 1 h of incubation with 7 U · ml-1 of lysozyme (sigma). B) Heat and hydrogen peroxide resistance assay. Percentage of resistant spores exposed for 7.5, 15 and 30 minutes at 90°C (filled symbols) or with 5% H_2_O_2_ (vol/vol) (open symbols) are shown. C) Hydrophobicity assay. The percentage of hydrophobicity represents the proportion (×100) of spores in n-hexadecane after the separation of the spores into water and solvent phases. D) Clumping assay of spores produced at 25 (blue) and 42°C (orange). The percentage of OD580nm decrease of spores suspended in distilled water was monitored for 15 minutes.

### 3.5 Spore morphology

Spore morphology appeared dependent on the growth temperature. Spores produced at the sub-optimal temperature (25°C) observed at the phase contrast microscope showed a regular and organized distribution (Fig. 7). All spores appeared phase bright, of elliptical shape and organized into shoulder-to-shoulder filaments (Fig. 7). Such organization was not observed when spores were produced at 42°C (Fig. 7). Since growing and sporulating cells produce more biofilm at 25 than at 42°C (Fig. 4), such organized distribution of spores might depend on cells entering into sporulation while entrapped in the extracellular matrix. The released spores would then keep the organized distribution of the sporulating cells.

**Fig. 7:**
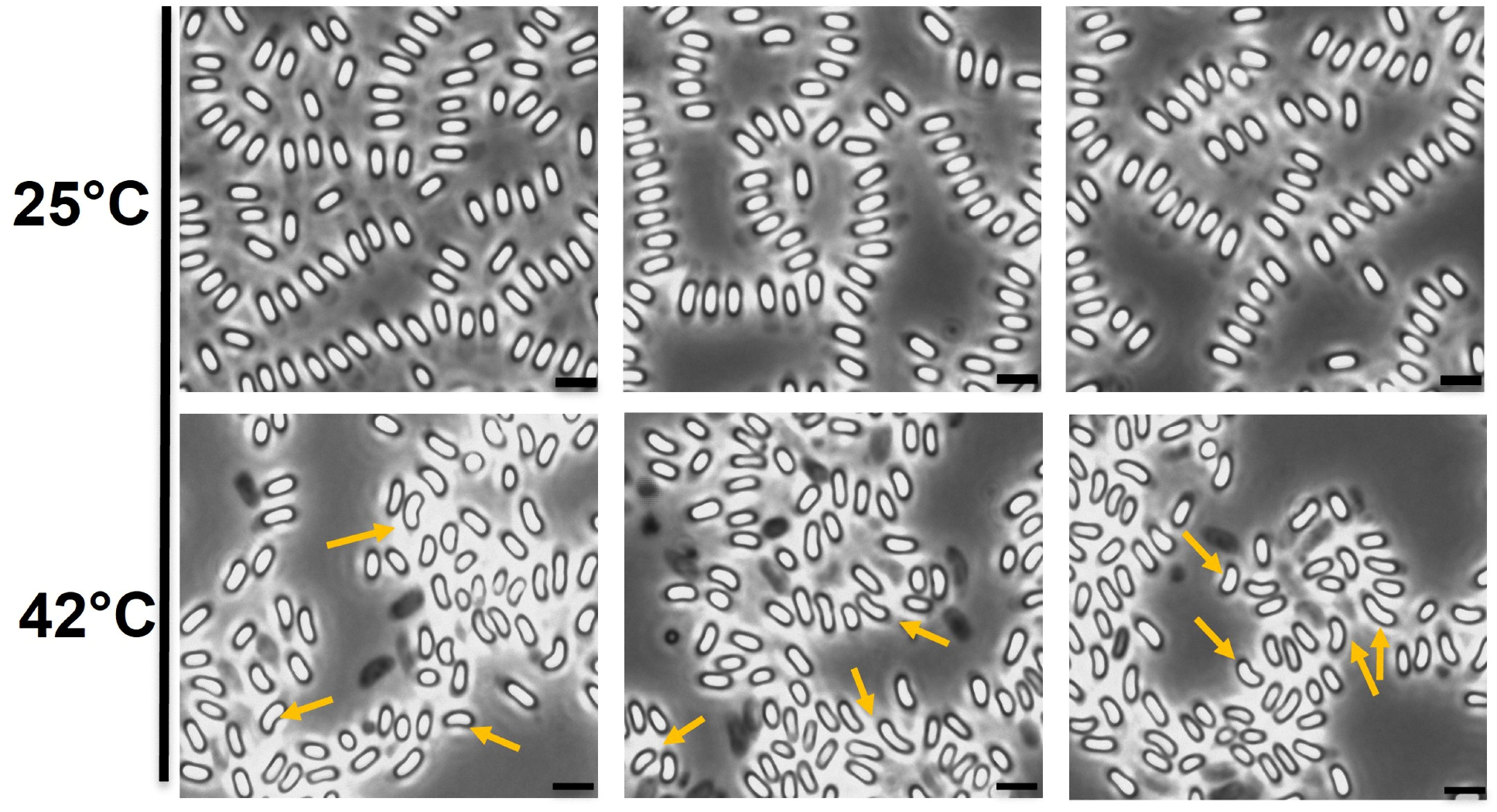
Morphology of spores produced at 25 and 42°C observed by phase-contrast microscopy. Arrows indicate the curved spores observed at the 42°C. For all panels the scale bar corresponds to 1μm.

Some of the spores produced at 42°C showed an unusual curved morphology (see Fig. 3CD and yellow arrows in Fig. 7), that was never observed in 25°C spores. The unusual morphology was strictly dependent on the growth temperature with no curved spores at 25°C and increasing numbers of curved spores raising the growth temperature from 37 (not shown) to 42°C. Counting over 300 spores in different microscopy fields, at 42°C about 30% of the spores showed the unusual morphology.

To further characterize the unusual morphology, purified spores were used for scanning electron microscopy analysis. As shown in Fig. 8, 25°C but not 42°C spores were enveloped into a matrix, supporting the hypothesis that such a matrix could be responsible for the organized spore distribution at 25°C observed in Fig. 7. In addition, the surface of spores appeared smooth at 25°C and wringled at 42°C (Fig. 8), suggesting that the wringled surface may be linked to the curved morphology.

**Fig. 8:**
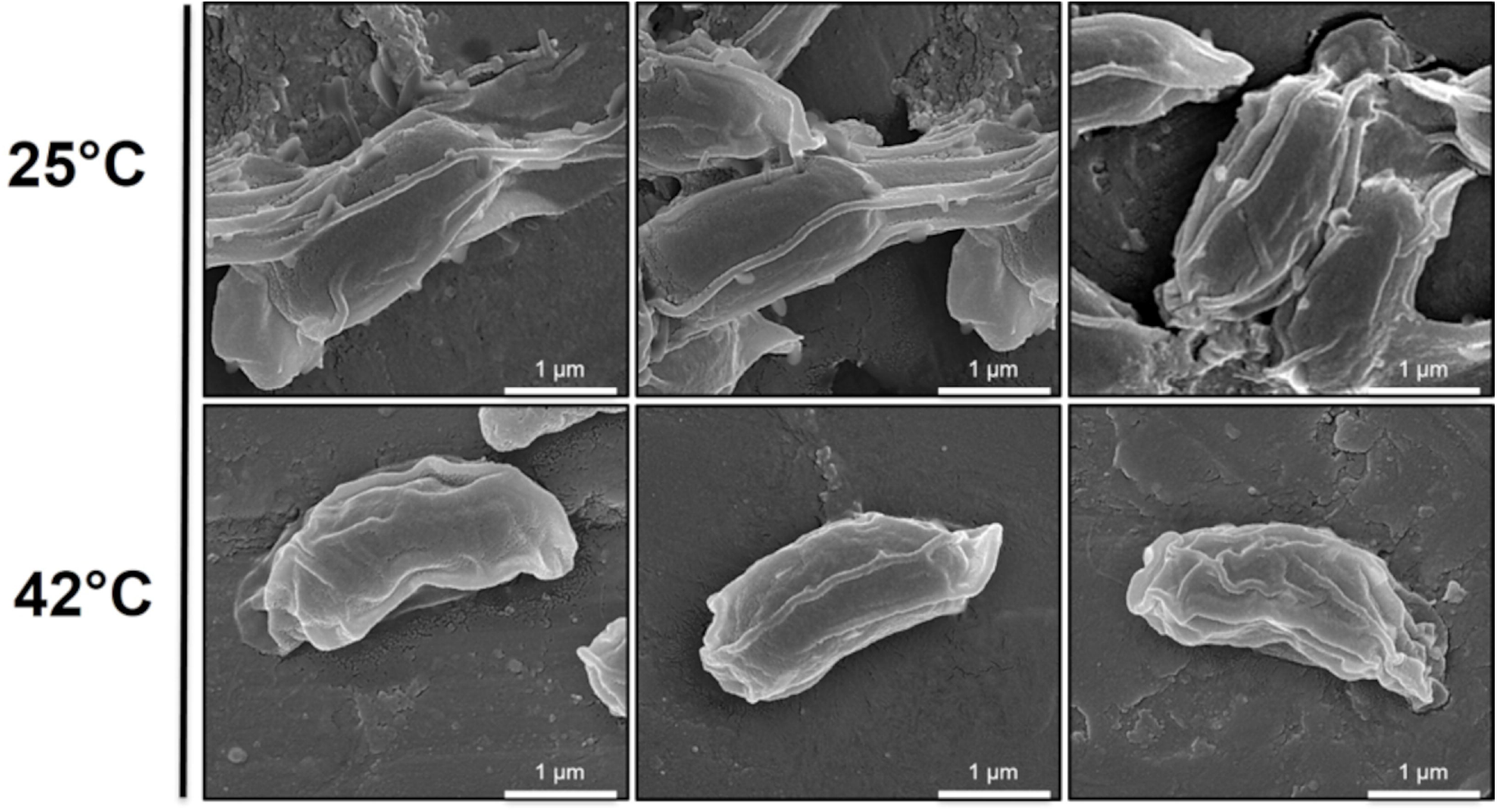
Scanning electron microscopy analysis of spores produced at the two different temperatures (25 and 42°C). Scale bar 1μm.

The unusual spore morphology observed at 42°C could be dependent on the assembly of the spore coat and exosporium proteins. The MV19 genome was analyzed for the presence of homologs of coat/exosporium genes by using as a “query” the list of the spore coat and exosporium proteins identified in the *B. cereus* reference strain ATCC 14579 (Abhyankar *et al.* 2013). As reported in the Suppl. Mat. Table S4, the majority of the putative MV19 proteins showed a high similarity with homologs of the type strain *B. cereus* ATCC 14579. Six proteins of the reference strain (BC_0944, BC_2426, BC_2427, BC_2569, BC_3515, BC_p0002) were not found in MV19 (Suppl. Mat. Table S4). These proteins are not characterized with the exception of BC_2569 annotated as “collagen triple helix repeats” and of BC_p0002 located on a plasmid of strain ATCC 14579.

## 4. Conclusions

MV19 is a human intestinal isolated strain belonging to the *B. cereus sensu stricto* species. Its genome contains genes coding for most known toxins and virulence-related factors of *B. cereus* with the exception of the genes required for Cereulide synthesis.

MV19 cells grow within a temperature range of 20 and 44°C with optimal growth at 37 and 42°C. At the optimal growth temperature of 42°C also spore production is faster than at the sub-optimal temperature of 25°C. However, the slow and prolonged growth at 25°C delayed the entry into sporulation allowing more cell divisions and the production of over 15-fold more free spores than at 42°C. The 25°C spores are also more resistant (to lysozyme, H_2_O_2_ and heat), more efficient in germination and more hydrophobic than those produced at 42°C. In addition, about 30% of the spores produced by fast growing cells showed an unusual curved morphology and a wringled surface, suggesting a defective structure that presumably affects the spore traits. Therefore, although at the optimal growth temperature of 42°C spore formation occurs faster and with a high efficiency, some of the produced spores are structurally and functionally defective. This conclusion suggests that the reduced rapidity and efficiency of sporulation at 25°C are compensated by a high quality and quantity of the released spores.

In *B. subtilis* a delay in spore formation negatively affects spore germination but increases the amount of spores produced and a quality-quantity trade-off has been proposed to occur and contribute to the evolutionary adaptation of spore formers (Mutlu et al., 2018; Mutlu et al., 2020). In *B. cereus* MV19 at the sub-optimal temperature of growth and sporulation more cells and more fully functional spores are produced than at the optimal growth temperature, compensating the faster growth and efficient sporulation observed at 42°C.

Although the molecular mechanisms regulating such balancing are totally obscure, some spore quality determinants have been proposed in *B. subtilis*. Examples are the electrical polarization of the outer membrane of developing spores (Sirec et al., 2019), the enzyme alanine dehydrogenase (Mutlu et al., 2018) and the isoleucyl-tRNA synthase (Kermgard et al., 2017) have all been proposed as quality control checkpoints. Homologs of both the alanine dehydrogenase (MV19_2153) and the isoleucyl-tRNA synthase (MV19_0519) are present in the MV19 genome and share, respectively, 72% and 73.86% of identity with the *B. subtilis* proteins. Addressing the role of these spore quality determinants in *B. cereus* would be a relevant future research challenge.

## Acknowledgements

The authors thank the electron microscopy facility (CESMA) of the Federico II University of Naples for their support with the SEM analysis.

## Author agreement

Authors approve the final version of the submitted manuscript.

## Authors’ contributions

AS, GDGB, SC and MV executed the experiments, AS, MV, SC, LB, RI and ER analyzed the data, interpreted the results. ER and AS wrote the manuscript, ER, RI and LB designed the study and provided the funding.

