## Supplementary Tables for "Sporulation efficiency and spore quality in a human intestinal isolate of *Bacillus cereus*"

**Table S1.** Toxin genes in the *Bacillus cereus* MV19 genome.

| ***Bacillus* toxins** | **Related genes** | ***Bacillus cereus* MV19^a,b^** |
| --- | --- | --- |
| **Anthrolysin O/Cereolysin O/Hemolysin** | *alo* | 99.80/100 |
| **Cereulide** | *cesA* | - |
|  | *cesB* | - |
|  | *cesC* | - |
|  | *cesD* | - |
|  | *cesH* | 100/100 |
|  | *cesP* | - |
|  | *cesT* |  |
| **Cytotoxin K** | *cytK* | 100/100 |
| **Hemolysin II** | *hlyII* | - |
| **Hemolysin III homolog** | *hlyIII-family* | 100/97 |
| **Hemolysin III** | *hlyIII* | 100/98 |
| **Hemolytic enterotoxin Hbl** | *hblA* | 97.64/100 |
|  | *hblC* | 100/99 |
|  | *hblD* | 99.75/100 |
| **Non-hemolytic enterotoxin Nhe** | *nheA* | 99.22/100 |
|  | *nheB* | 99.75/100 |
|  | *nheC* | 98.89/99 |

^a^ Values indicate the presence of the gene and represent the identity and coverage percentages for each virulence genes (%identity/%coverage) with respect to the reference strain ATCC 14579.

^b^ -: absence of the gene

**Table S2.** Virulence factors in the *Bacillus cereus* MV19 genome.

| **Class** | **Virulence factors** | **Related**  **genes** | ***B. cereus* MV19^a,b^** |
| --- | --- | --- | --- |
| **Enzyme** | Immune inhibitor A metalloproteinase | *inhA*^c^ | 100/100 |
|  |  | *inhA*^c^ | 99.75/100 |
|  | Phosphatidylcholine-preferring phospholipase C (PC-PLC) | *plcA* | 100/100 |
|  | Phosphatidylinositol-specific phospholipase C (PI-PLC) | *piplc* | 100/100 |
|  | Sphingomyelinase (SMase) | *sph* | 99.70/100 |
|  | Streptococcal enolase (*Streptococcus*) | *eno* | 100/100 |
| **Iron acquisition** | Bacillibactin | *dhbA* | 100/100 |
|  |  | *dhbB* | 98.65/100 |
|  |  | *dhbC* | 99.50/100 |
|  |  | *dhbE* | 97.40/100 |
|  |  | *dhbF* | 97.57/100 |
|  | IlsA | *ilsA* | 98.03/99 |
|  | Petrobactin | *asbA* | 99.34/100 |
|  |  | *asbB* | 99.18/99 |
|  |  | *asbC* | 100/100 |
|  |  | *asbD* | 98.72/100 |
|  |  | *asbE* | 99.06/100 |
|  |  | *asbF* | 99.29/100 |
| **Regulation** | PlcR-PapR quorum sensing |  |  |
|  |  | *papR* | 100/100 |
|  |  | *plcR* | 98.95/100 |

^a^ Values indicate the presence of the gene and represent the identity and coverage percentages for each virulence genes (% identity / % coverage)

^b^ -: absence of the gene

^c^ Two ORFs were assigned to the *inhA* gene

**Table S3**. Genes for biofilm synthesis in the MV19 genome

| **Gene ID** | **Description** | ***Bacillus cereus* MV19^a^** |
| --- | --- | --- |
| *purA* | adenylosuccinate synthetase | 99.7/100 |
| *purB* | adenylosuccinate lyase | 99.9/100 |
| *purC* | phosphoribosylaminoimidazolesuccinocarboxamide synthase | 99.7/100 |
| *purD* | phosphoribosylamine--glycine ligase | 98.5/100 |
| *purE* | phosphoribosylaminoimidazolesuccinocarboxamide synthase | 99.8/100 |
| *purF* | amidophosphoribosyltransferase | 99.8/100 |
| *purH* | bifunctional phosphoribosylaminoimidazolecarboxamide formyltransferase/IMP cyclohydrolase | 98.2/100 |
| *purK* | N5-carboxyaminoimidazole ribonucleotide synthase | 99.7/100 |
| *purL* | phosphoribosylformylglycinamidine synthase | 99/100 |
| *purM* | phosphoribosylformylglycinamidine cyclo-ligase | 99.6/100 |
| *purN* | phosphoribosylglycinamide formyltransferase | 98.8/100 |
| *purQ* | phosphoribosylformylglycinamidine synthase | 99.5/100 |
| *purS* | phosphoribosylformylglycinamidine synthase | 99.5/100 |
| *yezC* | putative transcriptional regulator | 100/100 |
| *sinR* | master regulator of biofilm formation | 96/100 |
| *sigB* | RNA polymerase sigma factor SigB | 95.2/100 |
| *epsA* | modulator of protein tyrosine kinase EpsB | 96.2/100 |
| *epsB* | [protein tyrosine kinase involved in biofilm matrix formation](https://www.ncbi.nlm.nih.gov/gene/938640) | 96.7/100 |
| *epsD* | [putative extracellular matrix glycosyltransferase](https://www.ncbi.nlm.nih.gov/gene/938611) | 96.1/100 |
| *epsG* | [biofilm extracellular matrix formation chain-length determining factor](https://www.ncbi.nlm.nih.gov/gene/937071) | 96/100 |
| *epsK* | [putative extracellular matrix component exporter; putative cyclic di-GMP receptor](https://www.ncbi.nlm.nih.gov/gene/8302980) | 96.7/100 |
| *epsM* | [putative O-acetyltransferase involved in biofilm matrix formation](https://www.ncbi.nlm.nih.gov/gene/936372) | 97/100 |
| *epsO* | pyruvyl transferase for matrix biofilm formation | 97/100 |
| *tapA* | [lipoprotein for biofilm formation](https://www.ncbi.nlm.nih.gov/gene/938532) | 98/100 |

**^a^** Values indicate the presence of the gene and represent the identity and coverage percentages (% identity / % coverage)

**Table S4.** Spore Coat and Exosporium Proteins encoded by the MV19 genome

| **Protein name**  **(*B. cereus* ATCC 14579)** | **Gene ID**  **(*B. cereus* ATCC 14579)** | **Target protein**  **(*B. cereus* MV19)** | **Protein identity (%)**  **VS *B. cereus*** **MV19^a^** |
| --- | --- | --- | --- |
| Alr1 | BC_0264 | MV19_3315 | 100 |
| BaxpA | BC_2149 | MV19_3219 | 100 |
| BC_0263 | BC_0263 | MV19_5510 | 99.7 |
| BC_0337 | BC_0337 | MV19_2225 | 100 |
| BC_0825 | BC_0825 | MV19_2496 | 99.2 |
| BC_0944 | BC_0944 | - | - |
| BC_0987 | BC_0987 | MV19_5523 | 100 |
| BC_0987 | BC_0987 | MV19_5523 | 100 |
| BC_0996 | BC_0996 | MV19_5716 | 99.3 |
| BC_1029 | BC_1029 | MV19_5164 | 100 |
| BC_1334 | BC_1334 | MV19_0978 | 100 |
| BC_1424 | BC_1424 | MV19_5282 | 99.8 |
| BC_1456 | BC_1456 | MV19_1778 | 98.6 |
| BC_1591 | BC_1591 | MV19_1644 | 99.8 |
| BC_1613 | BC_1613 | MV19_1621 | 99.3 |
| BC_1708 | BC_1708 | MV19_3847 | 100 |
| BC_2207 | BC_2207 | MV19_1978 | 98.8 |
| BC_2237 | BC_2237 | MV19_1948 | 99.8 |
| BC_2266 | BC_2266 | MV19_1919 | 98.9 |
| BC_2267 | BC_2267 | MV19_1918 | 100 |
| BC_2375 | BC_2375 | MV19_1813 | 100 |
| BC_2426 | BC_2426 | - | - |
| BC_2427 | BC_2427 | - | - |
| BC_2481 | BC_2481 | MV19_4246 | 99.1 |
| BC_2569 | BC_2569 | - | - |
| BC_2639 | BC_2639 | MV19_4465 | 94.1 |
| BC_2745 | BC_2745 | MV19_4134 | 99.7 |
| BC_2858 | BC_2858 | MV19_4961 | 98.9 |
| BC_2878 | BC_2878 | MV19_5459 | 98.5 |
| BC_2969 | BC_2969 | MV19_5056 | 99.1 |
| BC_3090 | BC_3090 | MV19_4587 | 99.4 |
| BC_3133 | BC_3133 | MV19_4506 | 98.7 |
| BC_3195 | BC_3195 | MV19_3951 | 97.7 |
| BC_3345 | BC_3345 | MV19_2757 | 97 |
| BC_3515 | BC_3515 | - | - |
| BC_3547 | BC_3547 | MV19_3675 | 98 |
| YodI | BC_3582 | MV19_3075 | 99.2 |
| BC_3784 | BC_3784 | MV19_0627 | 100 |
| BC_3786 | BC_3786 | MV19_0625 | 100 |
| BC_3787 | BC_3787 | MV19_0624 | 99.8 |
| BC_3986 | BC_3986 | MV19_0426 | 100 |
| BC_3992 | BC_3992 | MV19_0420 | 100 |
| BC_4387 | BC_4387 | MV19_0022 | 97.6 |
| BC_5056 | BC_5056 | MV19_3637 | 100 |
| BC_5181 | BC_5181 | MV19_3782 | 100 |
| BC_p0002 | BC_p0002 | - | - |
| BclB | BC_2382 | MV19_5786 | 93.6 |
| BclC | BC_3712 | MV19_3209 | 83.1 |
| BxpB | BC_1221 | MV19_0879 | 98.2 |
| CalY | BC_1281 | MV19_0925 | 100 |
| CotB1 | BC_0389 | MV19_2276 | 98.8 |
| CotB2 | BC_0390 | MV19_2277 | 99.3 |
| CotD | BC_1560 | MV19_1677 | 100 |
| CotE | BC_3770 | MV19_0641 | 100 |
| CotG | BC_2030 | MV19_3351 | 97.9 |
| CotJA | BC_0823 | MV19_2494 | 100 |
| CotJB | BC_0822 | MV19_2493 | 98.9 |
| CotJC | BC_0821 | MV19_2492 | 100 |
| CotN | BC_1279 | MV19_0923 | 99 |
| CotX1 | BC_2872 | MV19_5454 | 100 |
| CotX2 | BC_2874 | MV19_5456 | 95.9 |
| CotY | BC_1222 | MV19_0880 | 93 |
| Cotα | BC_4047 | MV19_0365 | 100 |
| CwlJ | BC_5390 | MV19_1399 | 97.1 |
| DacF | BC_4075 | MV19_0335 | 99.2 |
| Eno | BC_5135 | MV19_3824 | 100 |
| ExsFB | BC_2374 | MV19_1814 | 95.2 |
| ExsK | BC_2493 | MV19_4255 | 99.2 |
| ExsY | BC_1218 | MV19_0877 | 93.4 |
| GerQ | BC_5391 | MV19_1398 | 100 |
| GpR | BC_4319 | MV19_0092 | 98.1 |
| BC_1245 | BC_1245 | MV19_0903 | 79.7 |
| InA | BC_1284 | MV19_0928 | 99.9 |
| IunH | BC_2889 | MV19_5472 | 99.7 |
| IunH2 | BC_3552 | MV19_3670 | 100 |
| BC_2677 | BC_2677 | MV19_5399 | 99.3 |
| BC_0395 | BC_0395 | MV19_2282 | 99.5 |
| OppA | BC_2026 | MV19_3355 | 99.6 |
| RocA | BC_0344 | MV19_2231 | 100 |
| SafA | BC_4420 | MV19_1044 | 87.9 |
| SleB | BC_2753 | MV19_4126 | 99.6 |
| SodF | BC_1468 | MV19_1766 | 99.1 |
| SpoIVA | BC_1509 | MV19_1725 | 99.8 |
| SpoVT | BC_0059 | MV19_4856 | 100 |
| BC_4639 | BC_4639 | MV19_1264 | 98.8 |
| Tig | BC_4480 | MV19_1089 | 100 |
| YaaH | BC_3607 | MV19_3100 | 99.8 |
| YabG | BC_0047 | MV19_4844 | 100 |
| YabP | BC_0063 | MV19_4860 | 99 |
| YajC | BC_4410 | MV19_1034 | 100 |
| YhcN/CoxA | BC_4419 | MV19_1043 | 99.5 |
| YisY | BC_4774 | MV19_2940 | 100 |
| YpeB | BC_2752 | MV19_4127 | 100 |
| YppG | BC_1559 | MV19_1678 | 99.5 |
| YqfX1 | BC_1391 | MV19_5249 | 97.6 |
| YqfX2 | BC_2099 | MV19_3269 | 100 |
| YtfJ/GerW | BC_4640 | MV19_1265 | 100 |
| YtfJ1 | BC_2095 | MV19_3273 | 99.3 |
| YtfJ2 | BC_4640 | MV19_1265 | 100 |
| YusW | BC_0212 | MV19_4399 | 99.4 |
| YxeE | BC_3534 | MV19_3687 | 99.1 |
| BC_1613 | BC_1613 | MV19_0436 | 99.6 |

**^a^** The symbol “-“  indicates the absence of the protein in *B. cereus* MV19
