## Supplementary figures and images for "Sporulation efficiency and spore quality in a human intestinal isolate of *Bacillus cereus*"

### Supplementary Figure 1

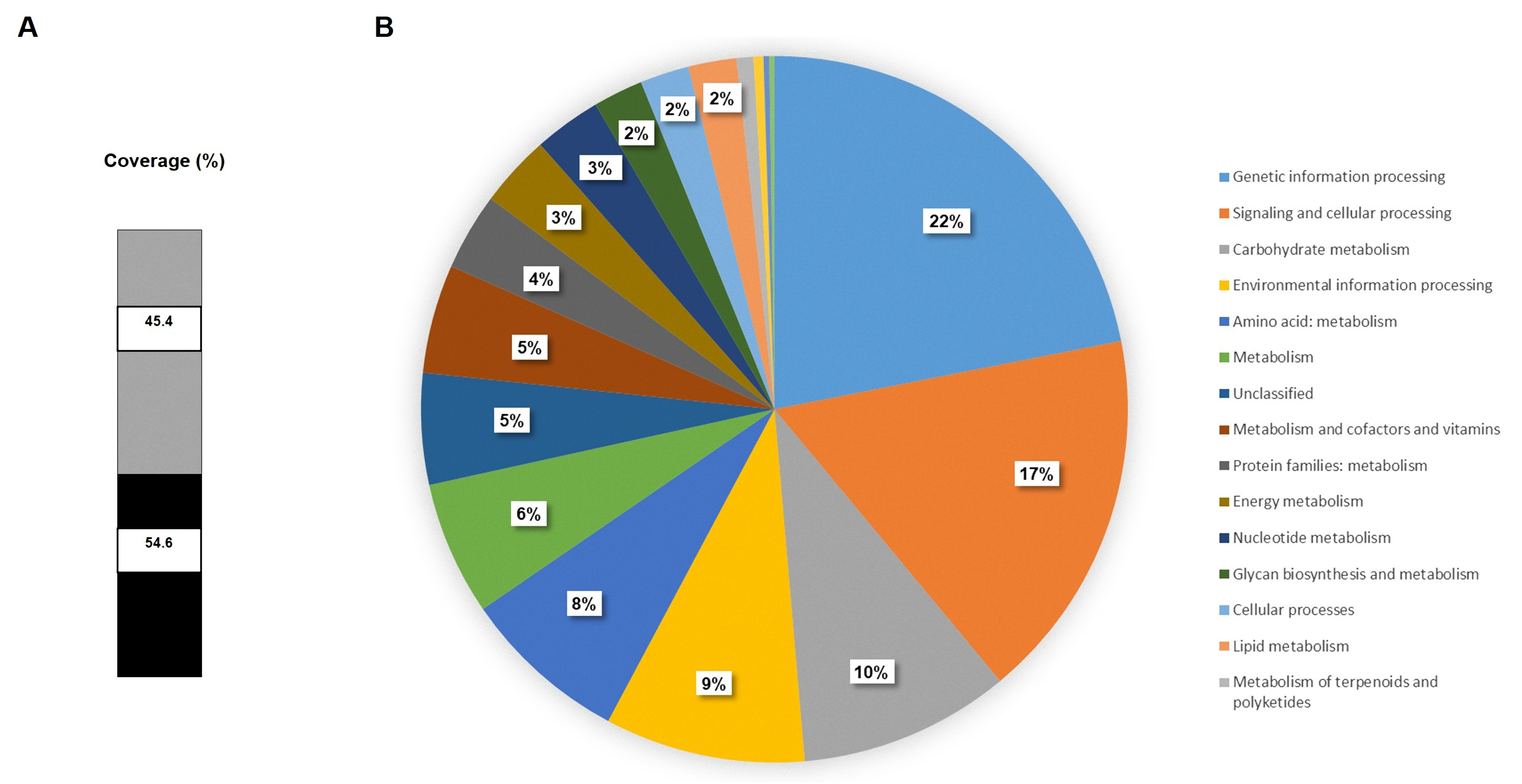

### Supplementary Figure 2

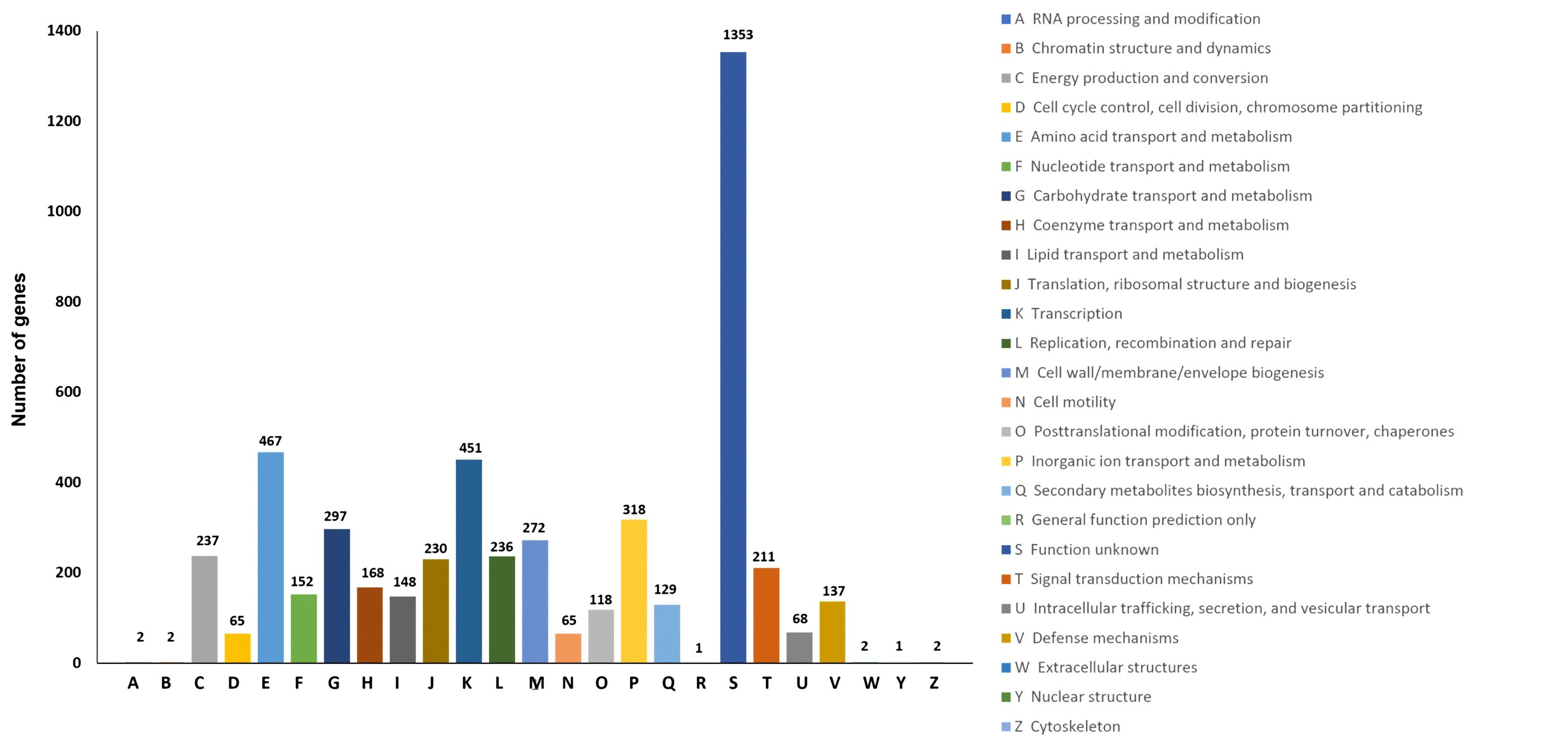

### Supplementary Figure 3

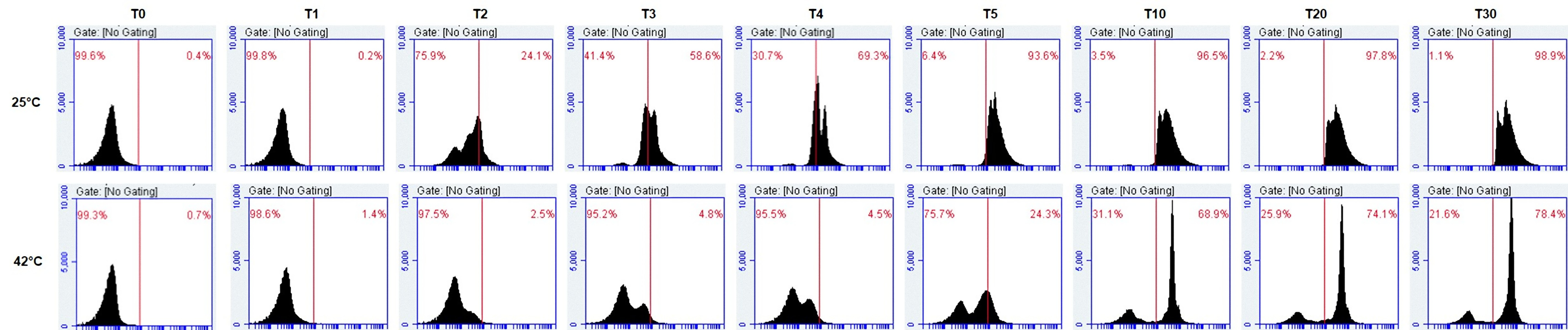
